# The glycosyltransferase UGT76B1 is critical for plant immunity as it governs the homeostasis of *N*-hydroxy-pipecolic acid

**DOI:** 10.1101/2020.06.30.179960

**Authors:** Lennart Mohnike, Dmitrij Rekhter, Weijie Huang, Kirstin Feussner, Hainan Tian, Cornelia Herrfurth, Yuelin Zhang, Ivo Feussner

## Abstract

The trade-off between growth and defense is a critical aspect of plant immunity. Therefore, plant immune response needs to be tightly regulated. The hormone regulating plant defense against biotrophic pathogens is salicylic acid (SA). Recently, *N*-hydroxy-pipecolic acid (NHP) was identified as second regulator for plant innate immunity and systemic acquired resistance. Although the biosynthetic pathway leading to NHP formation has already been identified, the route how NHP is further metabolized was unclear. Here, we present UGT76B1 as a UDP-dependent glycosyltransferase that modifies NHP by catalyzing the formation of 1-*O*-glucosyl-pipecolic acid (NHP-*O*Glc). Analysis of T-DNA and CRISPR knock-out mutant lines of *UGT76B1* by targeted and non-targeted UHPLC-HRMS underlined NHP and SA as endogenous substrates of this enzyme in response to *Pseudomonas* infection and UV treatment. UGT76B1 shows similar K_M_ for NHP and SA. *ugt76b1* mutant plants have a dwarf phenotype and a constitutive defense response which can be suppressed by loss of function of the NHP biosynthetic enzyme FMO1. This suggests that elevated accumulation of NHP contributes to the enhanced disease resistance in *ugt76b1*. Externally applied NHP can move to distal tissue in *ugt76b1* mutant plants. Although glycosylation is not required for the long distance movement of NHP during systemic acquired resistance, it is crucial to balance growth and defense.

## Introduction

Plants are constantly exposed to biotic and abiotic stress. To deal with external threats, plants have developed an impressive repertoire of chemical compounds. However, there is a trade-off between defense and growth as shown in autoimmune mutants such as *snc2*-1D *npr1*-1 and *s3h s5h*, which accumulate high levels of defense hormones and exhibit severe dwarf phenotypes (Zhang et al., 2010; Zhang et al., 2017). In order to balance between growth and defense, plants oversee the homeostasis of these compounds constantly. Dynamic changes of the levels of immune signaling molecules allow plants to react rapidly and appropriately to danger (Hartmann and Zeier, 2019; Huang et al., 2020). The biosynthesis, transport, and homeostasis of the signaling molecules is therefore, strictly regulated to prevent unintended consequences.

Two signaling molecules, salicylic acid (SA) and *N*-hydroxy-pipecolic acid (NHP), are particularly important in plant defense against biotrophic pathogens. Together they orchestrate the immune response in the local tissue to prevent pathogen spread (Hartmann et al., 2018; Guerra et al., 2020). Locally produced defense signals are further translocated to distal parts of the plant, leading to massive transcriptional and metabolic reprogramming in the naïve tissues, which enables a quick and robust response to subsequent infections (Bernsdorff et al., 2016). This induced immunity in distal tissue is termed systemic acquired resistance (SAR). Most of the signaling molecules participating in the induction of SAR can be found in the phloem upon infection (Fu and Dong, 2013). The effect of SA and NHP in the context of plant immunity has been well documented (Chen et al., 2018; Hartmann et al., 2018; Zhang and Li, 2019; Huang et al., 2020).

Biosynthesis of SA is divided into two major routes that result in SA formation in planta: The phenylpropanoid or PHENYLAMMONIA LYASE (PAL)-pathway and the ISOCHORISMIC ACID SYNTHASE 1 (ICS1)-pathway (Yalpani et al., 1993; Wildermuth et al., 2001). Nevertheless, in *Arabidopsis* about 90% of endogenous SA derives from chloroplast-derived isochorismic acid, which is exported to the cytosol via ENHANCED DISEASE SUSCEPTIBILITY 5 (EDS5) and conjugated to glutamate by AvrPphB SUSCEPTIBLE 3 (PBS3). The formed isochorismic acid-9-glutamic acid then spontaneously decomposes into SA and enolpyruvyl-*N*-glutamic acid (Rekhter et al., 2019b). Furthermore, ENHANCED PSEUDOMONAS SUSCEPTIBILITY 1 (EPS1) has been shown to enhance SA formation from isochorismic acid-9-glutamic acid (Torrens-Spence et al., 2019).

NHP was recently discovered as a signaling compound for plant defense against biotrophic pathogens (Chen et al., 2018; Hartmann et al., 2018). So far, research has focused on the biosynthesis of NHP from lysine. In the first step, the α-aminotransferase AGD2-LIKE DEFENSE RESPONSE PROTEIN 1 (ALD1) catalyzes the transamination of lysine into ε-amino-α-keto caproic acid (Song et al., 2004; Navarova et al., 2012; Vogel-Adghough et al., 2013). This compound spontaneously cyclizes and thereby yields Δ^1^-piperideine-2-carboxylic acid (P2C). In a second step, the ketimine reductase SAR DEFICIENT 4 (SARD4) catalyzes the formation of pipecolic acid (Pip) from P2C (Ding et al., 2016; Hartmann et al., 2017). Pip requires *N*-hydroxylation in order to reach its full protective capacity. This activation is catalyzed by FLAVIN-DEPENDENT MONOOXYGENASE 1 (FMO1) (Chen et al., 2018; Hartmann et al., 2018).

One important strategy to maintain a preferred concentration of an active metabolite is chemical modification, which can change the bioavailability and activity of the compound. Different modifications of SA such as hydroxylation and methylation have been described (Song et al., 2009; Zhang et al., 2017). SA itself as well as its catabolites can be further xylosylated (addition of the pentose xylose) and glycosylated (addition of a hexose) (Song et al., 2008; Bartsch et al., 2010; Huang et al., 2018). The transfer of an activated sugar moiety onto a target molecule is predominantly catalyzed by the widespread enzyme family of uridine diphosphate (UDP)-DEPENDENT GLYCOSYL TRANSFERASES (UGTs). The closely related UGT74F1 and UGT74F2 catalyze the formation SA-glycoside (SAG) and SA glucose ester (SGE) respectively (Dean and Delaney, 2008; George Thompson et al., 2017). Another enzyme UGT71C3 was recently shown to be responsible for the biosynthesis of methyl-SA glycoside (Chen et al., 2019). Despite the high abundance of these glycosides upon stress, the biological significance of the formation of these compounds is still elusive. Blocking glycosylation of SA has been shown to result in enhanced disease resistance (Noutoshi et al., 2012). In tobacco SAG is transported from the cytosol into vacuoles, suggesting that the glucosides are a storage form of SA. On the other hand, the formation of SAG may be important for the vascular transport, as there is evidence that SAG can be hydrolyzed back into SA in the extracellular space (Hennig et al., 1993; Seo et al., 1995).

So far, only one metabolite of NHP was identified, namely NHP-glycoside (NHP-*O*Glc) (Chen et al., 2018; Hartmann et al., 2018). Intriguingly, externally supplied NHP can be found in distal tissues in uninfected *fmo1* mutant plants as NHP and NHP-*O*Glc, suggesting that at least one of these molecules is mobile *in planta* (Chen et al., 2018). Currently, neither the function of NHP-*O*Glc nor the enzyme that catalyzes the glycosylation of NHP was identified. Here we report that UGT76B1, which was previously reported to glycosylate SA and 2-hydroxy-3-methyl-pentanoic acid (isoleucic acid, ILA), catalyzes the formation of NHP-*O*Glc (von Saint Paul et al., 2011; Noutoshi et al., 2012; Maksym et al., 2018). UGT76B1 has strong *in vitro* activity towards NHP and no detectable amount of NHP-*O*Glc is synthesized in *ugt76b1* mutant plants, which results in increased NHP accumulation, a dwarf phenotype and enhanced disease resistance against biotrophic pathogens. Moreover, we show that externally applied NHP is mobile to distal tissue in the absence of UGT76B1 and that transport of NHP seems not to depend on further glycosylation.

## Results

### Non-targeted metabolome analysis of infected leaf tissue revealed NHP as *in vivo* substrate of UGT76B1

Searching for the protein that catalyzes the formation of NHP-*O*Glc, we found *UGT76B1* as a recurring candidate gene in several studies (von Saint Paul et al., 2011; Noutoshi et al., 2012; Gruner et al., 2013; Hartmann et al., 2018). The loss-of-function mutant *ugt76b1-*1 showed enhanced resistance against *Pseudomonas* infections (von Saint Paul et al., 2011; Noutoshi et al., 2012; Maksym et al., 2018). Although UGT76b1 has previously been shown to exhibit SA glycosyltransferase activity, the enzyme has a high level of substrate promiscuity *in vitro*. Additional substrates are ILA, leucic acid, 2-ethyl-2-hydroxybutyric acid and valic acid (von Saint Paul et al., 2011; Noutoshi et al., 2012; Maksym et al., 2018). Since UGT76B1 has been shown to influence SA metabolism, we wondered if UGT76B1 has other substrates *in vivo.*

We conducted a non-targeted metabolome analysis on Col-0 and *ugt76b1*-1 leaves after mock or *Pseudomonas* treatment. The dataset obtained by the non-targeted UPLC-HRMS analysis contains 448 metabolite features (false discovery rate (FDR) < 0.005), which were arranged into 7 cluster by means of one-dimensional self-organizing maps. NHP-*O*Glc was not detectable in infected *ugt76b1*-1 mutant plants and SAG was strongly reduced compared to the *P.s.m.* infected wild type plants (Col-0; Figure 1, cluster 1). In contrast to that, NHP and SA showed a three-respective two-fold accumulation in infected *ugt76b1*-1 plants compared to the respective wild type plants (cluster 3). Interestingly, the NHP precursor Pip as well as 2HNG as fragment of the SA-precursor isochorismic acid-9-glutamic acid showed comparable amounts in infected wild type and *ugt76b1*-1 mutant plants (cluster 2). We could not find evidences for additional substrates or products of UGT76B1 under our conditions with the non-targeted approach. However, we detected increased levels of the second SA-derived metabolite SGE in *ugt76b1*-1 plants after infection (cluster 3). Together the experiment lead to the identification of NHP as *in vivo* substrate of UGT76B1.

**Figure 1.**
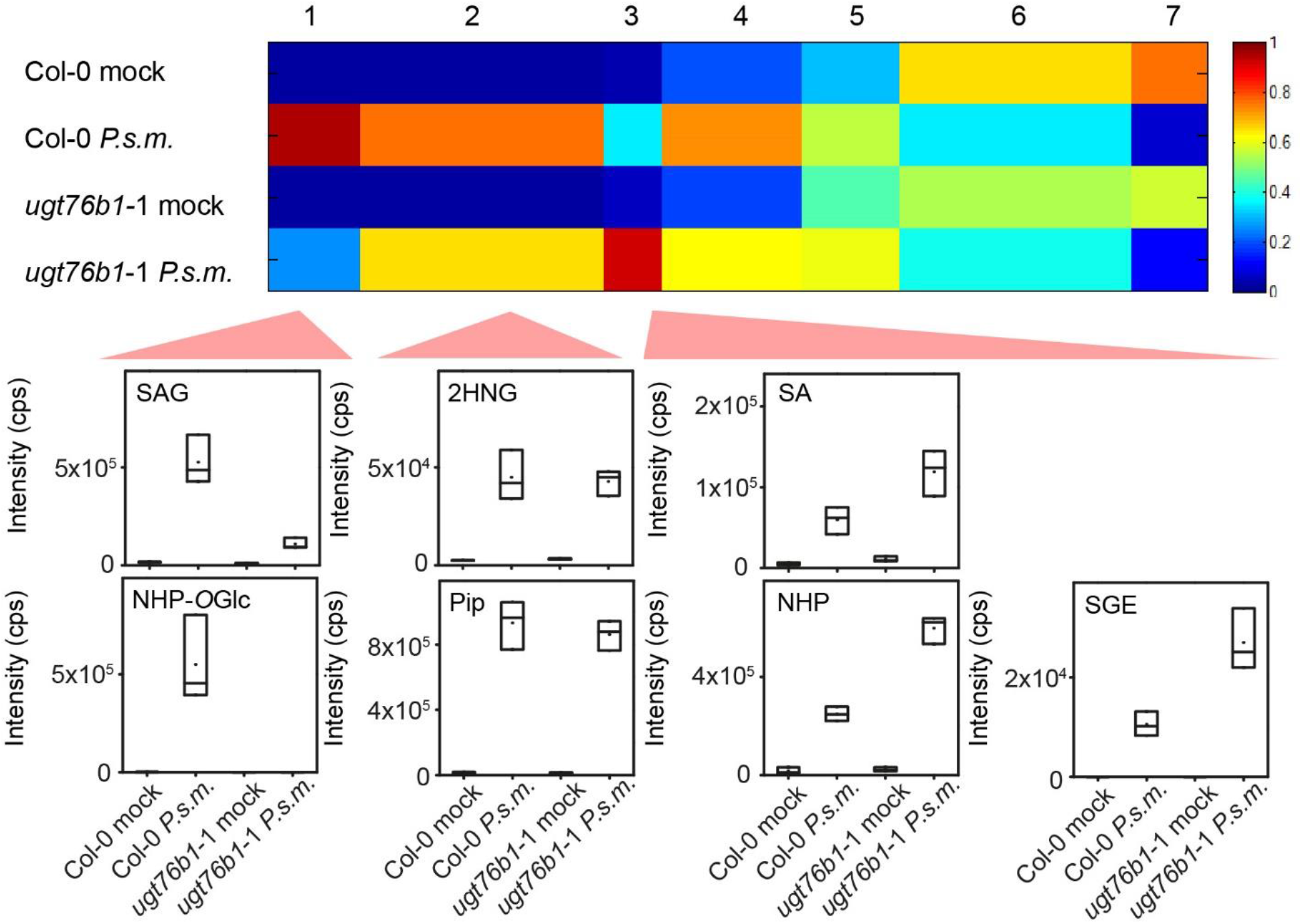
Non-targeted metabolomics revealed NHP as substrate of UGT76B1 *in vivo*. Col-0 and *ugt76b1*-1 mutant plants were infiltrated with MgCl_2_ (mock) or *Pseudomonas* ES4326 (*P.s.m.*) at OD_600_=0.05. Samples were collected 24 hours post infection. Metabolites of the polar extraction phase were analyzed by a metabolite fingerprinting approach based on UHPLC-HRMS. Intensity-based clustering by means of one-dimensional self-organizing maps of 448 metabolite features (FDR < 0.005) in 7 clusters is shown. The heat map colors represent average intensity values according to the color map on the right-hand side. The width of each cluster is proportional to the number of features assigned to this cluster. Box plots for selected metabolites of the indicated clusters are shown. The identity of the metabolites was unequivocally confirmed by UHPLC-HRMSMS analyses. The results were confirmed by two independent experiments. Data represents n=3 biological replicates.

### *UGT76B1* loss-of-function mutant plants do not accumulate NHP-*O*Glc

In addition to non-targeted metabolome analysis we quantitatively analyzed the amount of NHP, NHP-*O*Glc, SA and SAG in wild type (Col-0), *fmo1-*1 and *ugt76b1*-1 plants after infection with *Pseudomonas syringae* ES4326 (Figure 2a). 24 hours post infection (hpi), wild type plants accumulated NHP and NHP-*O*Glc to levels of 68 and 89 nmol/g FW, as well as of SA and SAG to 7 and 166 nmol /g FW, respectively. *ugt76b1*-1 plants exhibited a nearly three-fold higher accumulation of NHP (184 nmol/g FW) compared to wild type, whereas NHP-*O*Glc was not detected in the mutant after infection. As expected, *fmo1*-1 plants, which cannot generate NHP from Pip, accumulated neither NHP nor NHP-*O*Glc. Additionally, we observed an about 2.5-fold higher accumulation of SA after infection in *ugt76b1*-1 plants compared to wild type, whereas *fmo1*-1 plants exhibited comparable SA levels to the wild type, and moderately reduced SAG levels.

**Figure 2.**
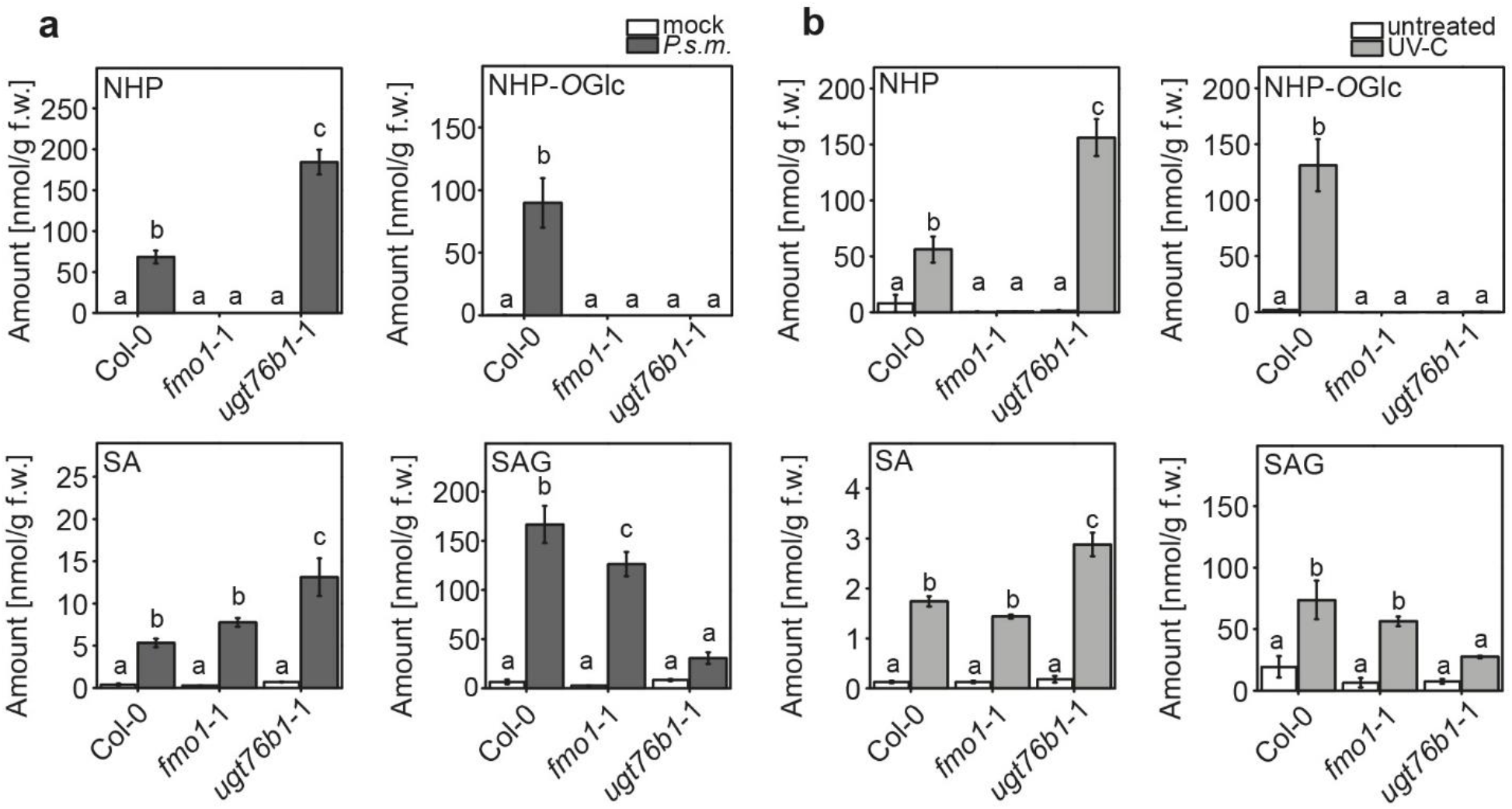
*UGT76B1* loss-of-function mutant plants are unable to synthesize NHP-*O*Glc. Amounts of *N*-hydroxy-pipecolic acid (NHP), NHP-glycoside (NHP-*O*Glc), salicylic acid (SA) and SA-glycoside (SAG) in wild type (Col-0), *fmo1*-1 and *ugt76b1*-1 plants after infection with *P.s.m.* ES4326 **(a)** or UV treatment **(b)**. Three leaves of 6-week-old plants, grown under short day conditions (8 hours light period), were infiltrated with *P.s.m.* ES4326 at OD_600_= 0.05 in 10 mM MgCl_2_ (*P.s.m.*) or 10 mM MgCl_2_ (mock). 24 hours post infiltration leaves were harvested and analyzed using UPLC-nanoESI-QTRAP-MS. Plants grown under long day conditions (16 h light period) were treated for 20 min with UV-C. 24 hours post UV-C treatment leaves were harvested and analyzed using UPLC-nanoESI-QTRAP-MS. Data represents the amount of analyte in nmol/g fresh weight (f.w.). Letters indicate statistical differences (p < 0.05, one-way ANOVA; n=3 biological replicates). The experiment was repeated once with similar results.

Similar results were obtained when we used UV-C to stimulate the production of NHP and SA independently of pathogen infection (Yalpani et al., 1994; Rekhter et al., 2019a). 24 h post UV-C-treatment, we detected 56 and 131 nmol/g FW of NHP and NHP-*O*Glc as well as 1.74 and 73 nmol/g FW of SA and SAG in wild type plants (Figure 2b). In *fmo1*-1 plants, no detectable amounts of NHP and NHP-*O*Glc were found after UV-C treatment, while SA and SAG accumulated to wild type levels. In *ugt76b1*-1 plants, we observed a nearly three-fold increase in NHP compared to wild type plants, but no formation of NHP-*O*Glc was detectable. There is also an increase in SA accumulation (2.87 nmol/g FW) and decrease in SAG accumulation (27 nmol/g FW) in *ugt76b1*-1. Together, these data strengthen the hypothesis that NHP-*O*Glc formation is dependent on a functional UGT76B1 enzyme, as additionally confirmed with two independent deletion mutant alleles of *UGT76B1* (Figure S1).

### UGT76B1 acts downstream of FMO1 thereby regulating plant immunity

We hypothesized that increased NHP accumulation in *ugt76b1*-1 plants after infection is due to its impaired glycosylation and that the dwarfed and enhanced resistance phenotype requires NHP. Furthermore, we assumed that UGT76B1 acts downstream of FMO1. To test this hypotheses, we checked growth of *Hyaloperonospora arabidopsis* (*H*. *a*.) Noco 2 on Col-0, *fmo1-*1, *FMO1*-3D (a gain-of-function mutant for *FMO1*), three mutant alleles of *UGT76B1* (*ugt76b1*-1, -3 and -4) and three *fmo1-*1 *ugt76b1* double knock-out mutant lines (*fmo1-*1 *ugt76b1-5*, *fmo1*-1 *ugt76b1*-1-40 and *fmo1*-1 *ugt76b1*-1-104; Figure 3). In comparison to Col-0, *FMO1*-3D showed high resistance against *H*. *a*. Noco 2, while *fmo1*-1 was more susceptible. *ugt76b1*-1, -3 and -4 exhibited strong resistance, but the double mutant lines showed similar susceptibility as *fmo1*-1 (Figure 3a). Additionally, we found that basal *PR1* gene expression is enhanced in all three *ugt76b1* alleles compared to Col-0 (Figure 3b), consistent with findings from a previous report (von Saint Paul et al., 2011). In contrast, the expression level of *PR1* is similar in *fmo1*-1 *ugt76b1*-5 and *fmo1*-1. In addition, the dwarf phenotype and dark green leaf color in the *ugt76b1* alleles are suppressed in the *fmo1*-1 *ugt76b1*-5 double mutant (Figure 3c). The *fmo1*-1 *ugt76b1*-1 double mutant plants accumulate neither NHP nor NHP-*O*Glc (Figure S2). Altogether, the data indicate that UGT76B1 acts downstream of FMO1 and that NHP is required for both the enhanced resistance and dwarf phenotype of *ugt76b1* plants.

**Figure 3.**
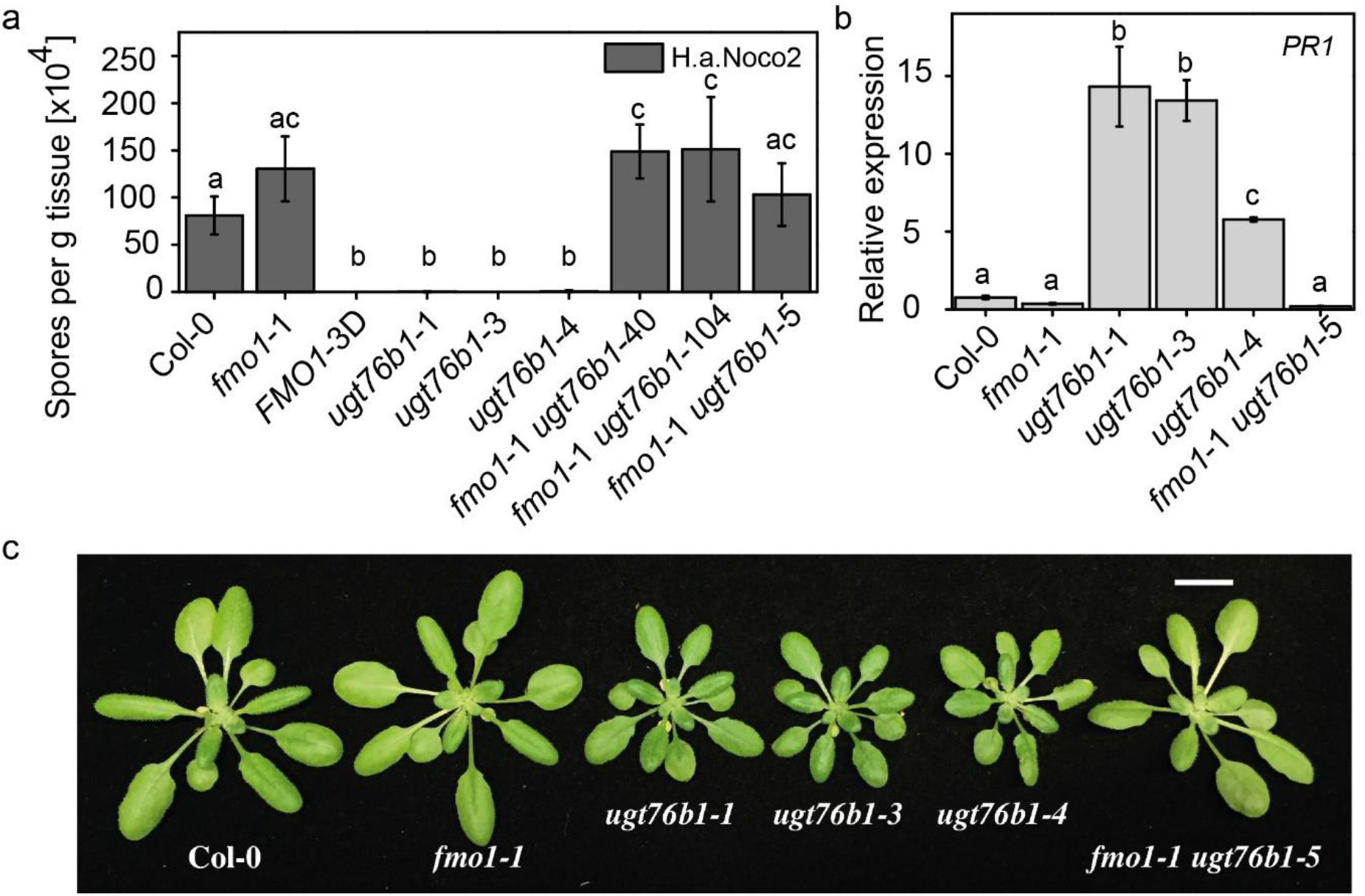
Rescue of *ugbt76b1* mutant phenotypes by introduction of the *fmo1-*1 mutation. **(a)** Growth of *H. a*. Noco2 on wild type (Col-0), *fmo1*-1, *FMO1*-3D, *ugt76b1*-1, *ugt76b1*-3, *ugt76b1*-4, *fmo1*-1 *ugt76b1*-40, *fmo1*-1 *ugt76b1*-104 and *fmo1*-1 *ugt76b1*-5 plants. Two-week-old seedlings were sprayed with *H.a.* Noco 2 spore suspension (5 × 10^4^ spores/mL). Infection was scored 7 days after infection. Letters indicate statistical differences (p < 0.05, one-way ANOVA; n=4 biological replicates). **(b)** Basal *PR1* gene expression in four-week-old plants of the indicated genotypes determined via quantitative RT-PCR. Letters indicate statistical differences (p < 0.05, one-way ANOVA; n=3 biological replicates). **(c)** Growth phenotypes of Col-0, *fmo1*-1, *ugt76b1*-1, *ugt76b1*-3, *ugt76b1*-4 and *fmo1*-1 *ugt76b1*-5. The Photo was taken on four-week-old plants grown under long day conditions (16 hours light/8 hours dark cycle). Scale bar is 1 cm.

### Increased accumulation of NHP in *ugt76b1* plants underlines the importance of turnover via UGT76B1

Next, we wondered whether the enhanced accumulation of NHP and SA in the *ugt76b1* mutants after infection is due to impaired turnover or increased biosynthesis of NHP and SA. Therefore, we measured the transcript levels of SA and NHP biosynthetic genes 24 hpi with Pseudomonas by quantitative RT-PCR. The transcript abundance of the SA biosynthetic genes *ICS1*, *EDS5* and *PBS3* (Figure 4a, 4b and 4c) was similar in the wild type and *ugt76b1-*1 mutant. Interestingly, transcripts of all three genes were upregulated in the mock-treated *ugt76b1-*1, suggesting that the basal levels of these SA biosynthetic genes are higher in the *UGT76B1* knock-out background. This is supported by the transcript levels of *PR1* and *PR2* after mock treatment (Figure S3). Despite the increased amount of NHP (Figure 2a), the transcript levels of NHP-biosynthetic genes *ALD1* and *FMO1* are significantly reduced in *ugt76b1*-1 compared to wild type. As a control, we monitored the transcript level of *UGT74F2* in Col-0 and *ugt76b1*-1. The transcript abundance of *UGT74F2* did not change after infection in Col-0 and *ugt76b1*-1 plants (Figure 4). Taken together, the increased SA and NHP levels in *ugt76b1* mutants upon pathogen infection is unlikely caused by their increased biosynthesis, since the respective transcripts are not higher in *ugt76b1*-1 than in wild type. These findings support that UGT76B1 plays a central role in the turnover of NHP and influences the formation of SAG.

**Figure 4.**
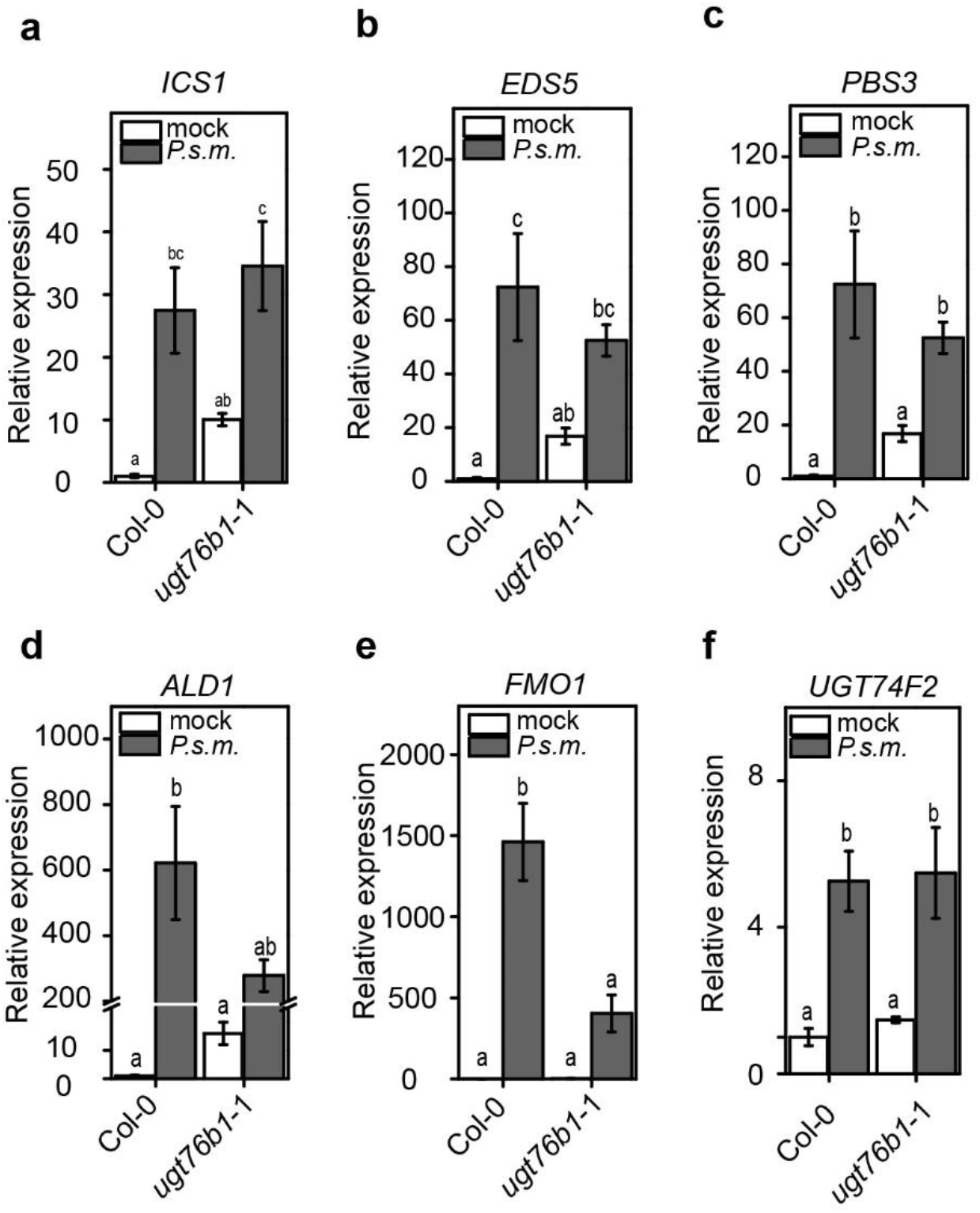
Comparisons between transcript levels of *ICS1*, *EDS5*, *PBS3*, *ALD1*, *FMO1* and *UGT74F2* in *ugt76b1* and wild type. Transcript abundance of genes encoding SA and NHP biosynthetic enzymes was analyzed in wild type and *ugt76b1*-1 plants after infection with *P.s.m.* ES4326. Three leaves of 4-6 week-old plants were treated with *P.s.m.* ES4326 (OD_600_=0.001). Leaves were harvested 24 hours post infection and analyzed via quantitative PCR using cDNA generated by reverse-transcriptase reaction as templates. Letters indicate statistical differences (p < 0.05, one-way ANOVA; n=3 biological replicates). Graph **d** includes an axis break from 25 to 200.

### UGT76B1 catalyzes the glycosylation of NHP *in vitro*

In addition, we checked whether UGT76B1 can glycosylate NHP *in vitro*. The His-tagged UGT76B1 was heterologously expressed in *Escherichia coli* and purified to homogeneity by affinity chromatography and size exclusion chromatography (Figure S4). The enzymatic reaction of recombinant UGT76B1 with NHP and UDP-glucose as substrates was monitored by ultra-high performance liquid chromatography coupled to high-resolution mass spectrometry (UHPLC-HRMS). As shown in Figure 5a, UGT76B1 catalyzes *in vitro* formation of NHP-*O*Glc (*m*/*z* 308.1342, retention time [RT] 2.12 min). We also confirmed glycosylation of SA and ILA by UGT76B1 (von Saint Paul et al., 2011; Noutoshi et al., 2012). The formation of the respective glucosides SAG (*m*/*z* 299.0793, RT 3.14 min) and ILA-glycoside (ILA-Glc) (*m*/*z* 293.1240, RT 3.35 min) is shown in Figure 5c and 5b. In addition, we determined the Michaelis-Menten constant (K_M_) for SA and NHP. We analyzed the respective product signal area for NHP-*O*Glc and SAG via UHPLC-HRMS, resulting in K_M_(NHP) = 86±7 μM and K_M_(SA) = 90±7 μM (Figure 5d and 5e), which suggest that UGT76B1 has similar affinity towards SA and NHP. Together, our *in vitro* analysis shows that the purified recombinant UGT76B1 was active with about 5-fold higher affinity towards NHP and SA in comparison to the substrate ILA (Maksym et al., 2018).

**Figure 5.**
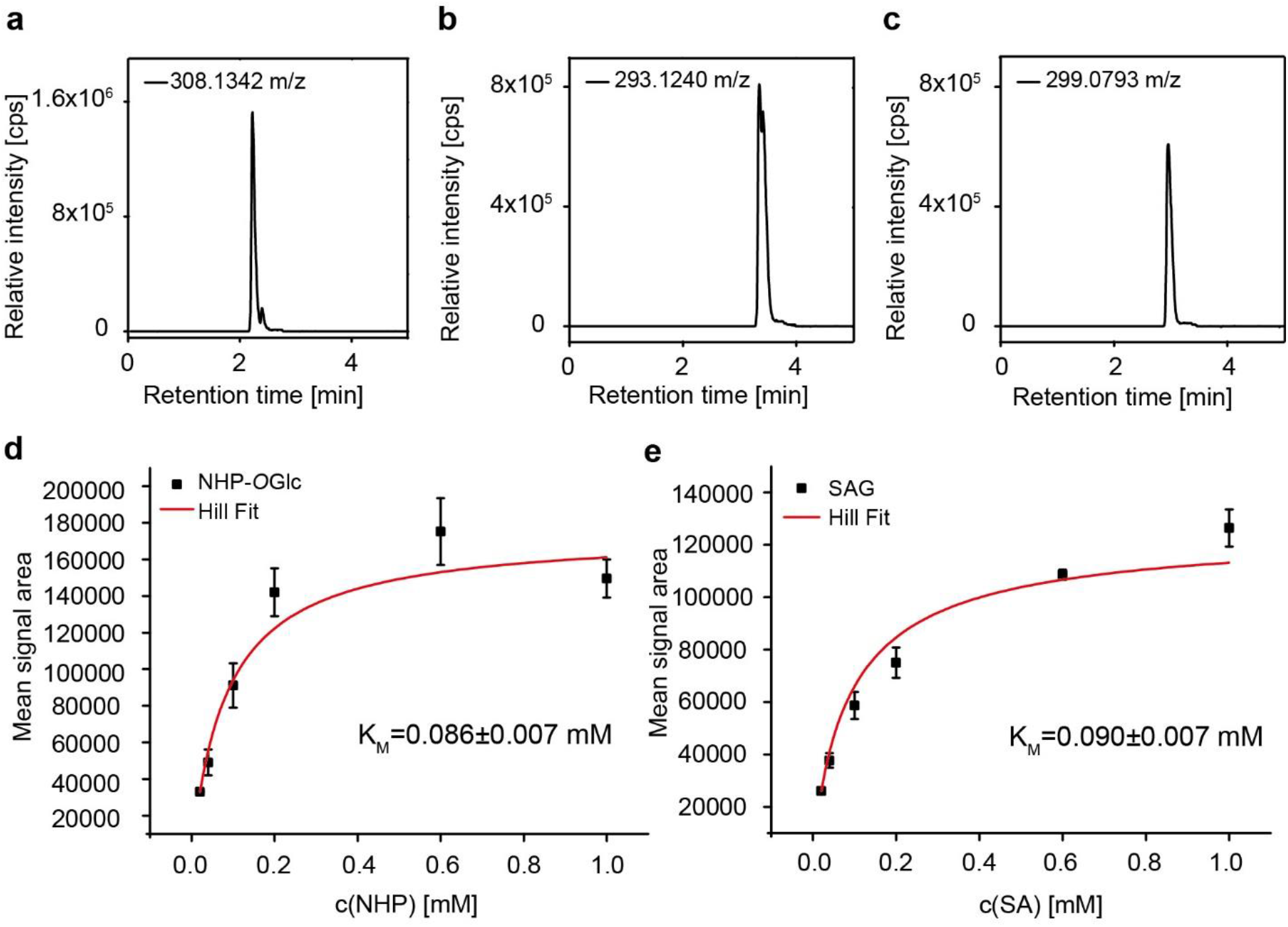
Glycosylation of SA, ILA and NHP by UGT76B1 *in vitro*. Activity assays were carried out using NHP, ILA and SA as substrates for the recombinant UGT76B1. Extracted ion chromatograms of the reaction products **(a)** NHP-*O*Glc (*m*/*z* 308.1342), **(b)** isoleucic acid-glycoside (ILA-Glc) (*m*/*z* 293.1240) and **(c)** SAG (*m*/*z* 299.0793) are shown. 10 μg of recombinant UGT76B1 were incubated with 50 μM substrate and 500 μM UDP-Glc at 30 °C for 30 min. The reaction was stopped by adding 25 % (*v/v*) acetonitrile. Michaelis-Menten constants (K_M_) of UGT76B1 were determined for the substrate NHP (Coefficient of determination (R^2^)=0.974) **(d)** and SA (R^2^=0.993) **(e)**, respectively. Mean signal area of the respective products (NHP-*O*Glc or SAG) from three replicates at different substrate concentrations are shown. Non-linear Hill regression was performed with Origin Pro 8.5 (OriginLab Corporation, Northhampton, MA, USA). All samples were measured via UHPLC-HRMS-analysis. Data are representative for two independent experiments.

We further analyzed active site residues in enzymes capable of glycosylating SA (UGT74F1 and UGT74F2) and compared them with the UGT76B1 protein sequence (Figure S5a). In addition, we made an *in silico* structural prediction of UGT76B1 using the deposited structure of UGT74F2 (PDB accession 5V2J, (George Thompson et al., 2017)) and modeled NHP in the electron density of the co-crystalized SA-analogue 2-bromobenzoic acid (Figure S5b and S5c). Some residues such as histidine at position 20 (His20) and aspartic acid at position 109 (Asp109) that have been shown to be important for the formation of SAG and SGE are conserved in all three UGTs (Figure S5a) (George Thompson et al., 2017). However, two threonine residues involved in the glycosylation of SA in UGT74F2 are substituted by leucine at position 17 (Leu17) and glycine at position 363 (Gly363) (Figure S5a and S5c). Nevertheless, we identified a threonine at position 131 in a predicted loop region, which might compensate the lack of Thr17 and Thr363 in the catalytic reaction (Figure S5a and S5c). These findings support our experimental data that the minimum subset of amino acids for fulfilling the glycosylation reactions on SA and NHP are present in UGT76B1’s putative active site.

### Deuterated NHP is translocated to distal tissue

NHP is the biological active metabolite of Pip in plant defense, especially in SAR (Chen et al., 2018; Hartmann et al., 2018). Nevertheless, it is still an open question whether NHP or NHP-*O*Glc might act as a mobile signal in SAR (Chen et al., 2018; Holmes et al., 2019). To address this question, we infiltrated uniformly deuterated NHP (D_9_-NHP) into leaves of Col-0, *fmo1*-1 and *ugt76b1*-1 plants. 24 hours post infiltration, local as well as systemic leaves were harvested. First, the formation of D_9_-NHP-*O*Glc that derived from the infiltrated D_9_-NHP in the local leaves of Col-0, *fmo1*-1 and *ugt76b1*-1 plants was monitored by UHPLC-HRMS. As expected, the applied D_9_-NHP was converted to D_9_-NHP-*O*Glc in the local leaves of wild type and *fmo1*-1 plants, but no D_9_-NHP-*O*Glc was detectable in *ugt76b1*-1 plants (Figure 6). Accordingly, the relative signal area of D_9_-NHP was two times higher in the local leaves of *ugt76b1*-1 plants in comparison to Col-0. Further analysis showed that D_9_-NHP was present in systemic tissue of the three genotypes Col-0, *fmo1*-1 and *ugt76b1*-1, whereas D_9_-NHP-*O*Glc was only detectable in Col-0 and *fmo1*-1 plants. This indicates that D_9_-NHP can move to distal tissues without glycosylation.

**Figure 6.**
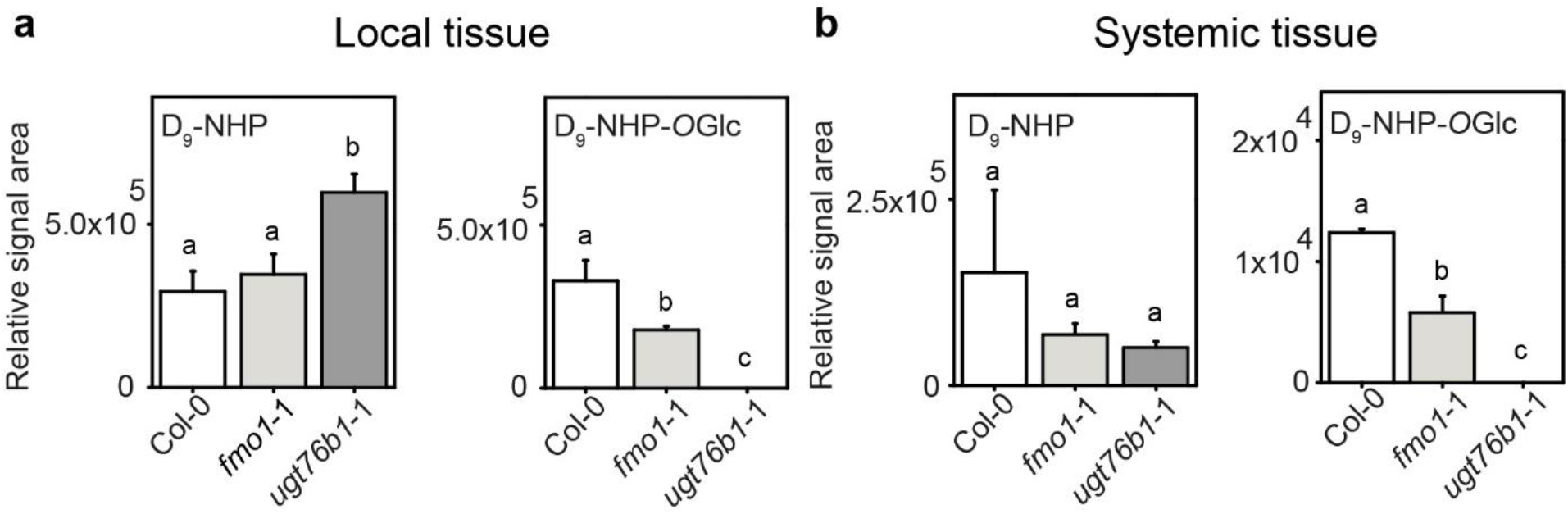
Infiltrated D_9_-NHP moves systemically and is converted to D_9_-NHP-OGlc in wild type and *fmo1*-1 but not in *ugt76b1*-1 plants. Relative intensities of deuterated NHP (D_9_-NHP) and its glucoside D_9_-NHP-*O*Glc were analyzed 24 hours after infiltration of D_9_-NHP to local tissue. Local and systemic leaves were harvested and analyzed by UHPLC-HRMS. Letters indicate statistical differences (p < 0.05, one-way ANOVA; n=3 biological replicates). The experiment was repeated once with similar results.

### *ugt76b1* plants exhibit enhanced resistance in systemic tissue

Next, we analyzed whether *ugt76b1*-1 can still establish SAR without the accumulation of NHP-*O*Glc by conducting a *H.a.* Noco 2 growth assay on plants pre-treated with *Pseudomonas syringae* (Figure 7). Establishment of SAR strongly reduces the disease rate of distal leaves (indicated as disease categories from 0 to 5) during a second infection with *H.a.* Noco 2, as shown for Col-0 plants (Figure 7a). Plants mock treated on the primary leaf showed high infection rates, indicated by disease categories of four and five on the systemic leaves after plant *H.a.* Noco 2 infection. For *ugt76b1*-1 plants, infection on the systemic leaves was reduced to minimum (disease category 0) no matter whether they were pre-induced with *Pseudomonas* or not. These disease rates were as low as those known for the *FMO1*-3D mutant. In contrast, *fmo1*-1 plants are not able to establish SAR and show therefore an increased susceptibility to *H.a.* Noco 2 as known from the literature (Ding et al., 2016). This finding indicates that the distal parts of *ugt76b1*-1, regardless of a primary infection, exhibit enhanced resistance towards *H.a.* Noco 2. This is consistent with results from our local *H.a.* Noco 2 infection assays for the *ugt76b1* lines (Figure 3a). In an independent approach, we analyzed the resistance of *ugt76b1*-1 to a secondary infection by *Pseudomonas*. As expected, Col-0 established SAR after primary infection, *fmo1*-1 plants were not able to establish SAR, and *FMO1*-3D showed a constitutive SAR phenotype (Figure 7b). Nevertheless, *ugt76b1*-1 exhibited reduced bacterial growth in distal leaves of both mock and *P.s.m.*-treated samples. Together, these data suggest that *ugt76b1*-1 displays constitutive resistance towards pathogens.

**Figure 7.**
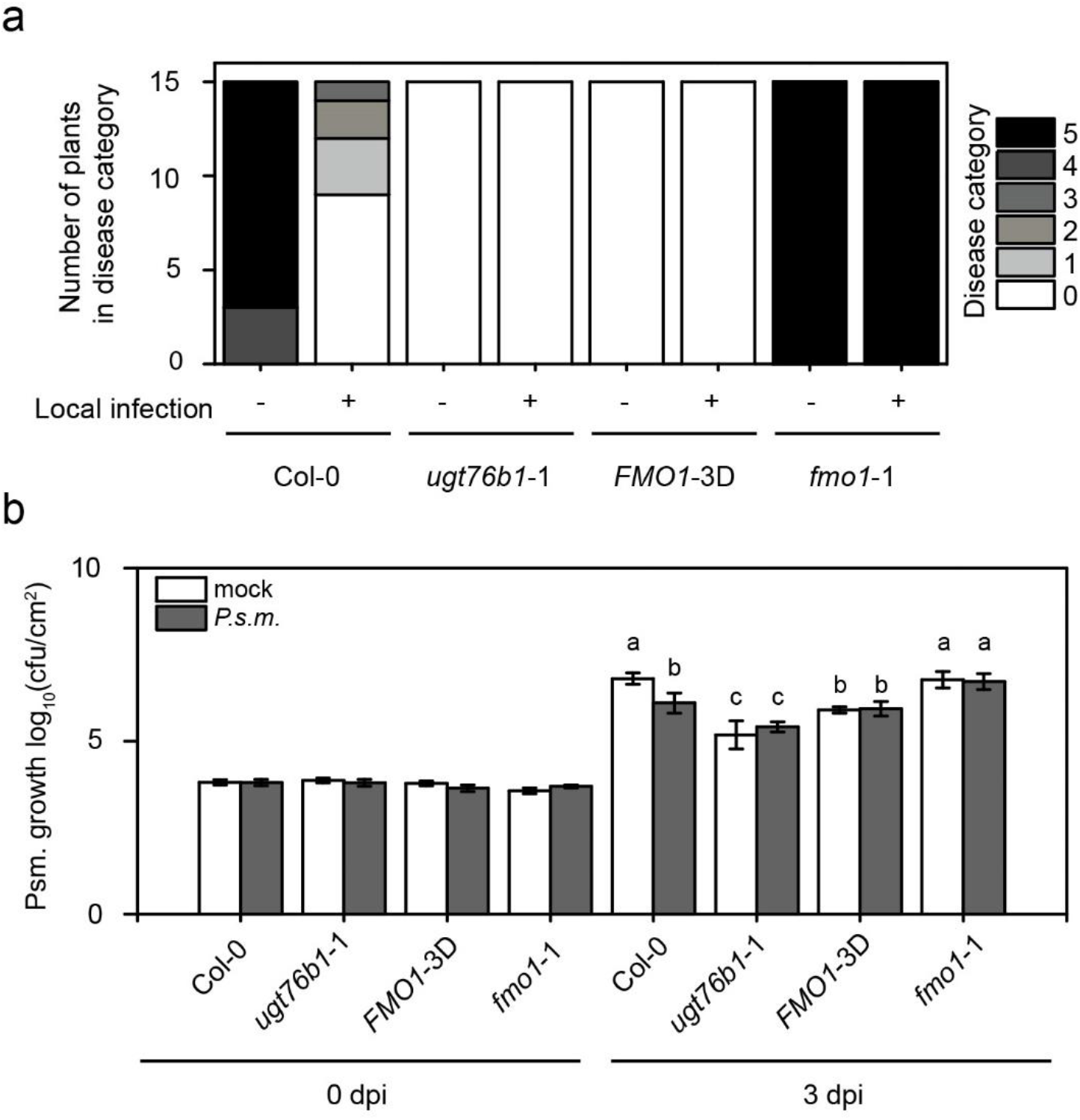
Growth of *H.a.* Noco2 and *P.s.m* on the distal leaves of wild type (Col-0), *ugt76b1*-1, *FMO1*-3D and *fmo1*-1. Three-week-old plants were first infiltrated with *P.s.m.* ES4326 (OD_600_ = 0.001) or 10 mM MgCl_2_ (mock) on two primary leaves and sprayed with *H. a.* Noco 2 spores (5 × 10^4^ spores/mL) 2 days later. Infections on systemic leaves were scored 7 days after inoculation as described previously (Zhang et al., 2010). A total of 15 plants were scored for each treatment. Disease rating scores are as follows: 0, no conidiophores on the plants; 1, one leaf was infected with no more than five conidiophores; 2, one leaf was infected with more than five conidiophores; 3, two leaves were infected but no more than five conidiophores on each infected leaf; 4, two leaves were infected with more than five conidiophores on each infected leaf; 5, more than two leaves were infected with more than five conidiophores. Similar results were obtained in three independent experiments **(a)**. Four-week-old plants were first infiltrated with *P.s.m.* ES4326 (OD_600_ = 0.001) or 10 mM MgCl_2_ (mock) on two primary leaves. Two days later, two upper leaves were challenged with *P.s.m.* ES4326 (OD_600_ = 0.001). Infections on systemic leaves were scored directly after (0 dpi) and three days post inoculation (3 dpi). Letters indicate statistical differences (p < 0.05, one-way ANOVA; n=6-8 biological replicates). Similar results were obtained in three independent experiments **(b)**.

## Discussion

The identification of FMO1 as a NHP biosynthetic enzyme was a major breakthrough towards the understanding of Pip-mediated plant immunity and its involvement in the establishment of SAR (Chen et al., 2018; Hartmann et al., 2018; Holmes et al., 2019). In addition, NHP-*O*Glc was recently described as metabolite of NHP (Chen et al., 2018). However, the enzyme catalyzing the formation of NHP-*O*Glc was unknown. In this study, we identified UGT76B1 as the enzyme responsible for the glycosylation of NHP *in vivo* and *in vitro* - in addition to its previously identified substrates SA and ILA. Beside its glycosyltransferase activity toward NHP *in vitro*, we show that UGT76B1 is required for the formation of NHP-*O*Glc *in planta* during pathogen infection. The absence of UGT76B1 leads to a significantly increased accumulation of NHP, the regulator of plant immunity, and the complete depletion of NHP-*O*Glc in *ugt76b1* mutant plants. Our data emphasize UGT76B1 as the only enzyme which glycosylates NHP *in planta*.

*ugt76b1* mutants have been shown to exhibit enhanced disease resistance against biotrophic pathogens, which was suggested to be caused by increased accumulation of SA (Noutoshi et al., 2012). The substrate ILA was recently suggested to activate immune response via SA by inactivating UGT76B1 (Bauer et al., 2020). In *ugt76b1* mutants, however, NHP accumulates to considerably higher level than in wild type during pathogen infection, suggesting that the elevated NHP level instead, may play a major role contributing to the enhanced disease resistance in the mutant plants. This is supported by the complete suppression of the autoimmune phenotype of *ugt76b1* by loss of function of FMO1. The accumulation of NHP leads to dwarfism as reported for the *FMO1*-3D overexpression line. Furthermore, increased NHP levels leads to enhanced resistance of this mutant (Koch et al., 2006). In contrast, the plant size increases if the amount of NHP decreases and its susceptibility towards biotrophic pathogen increases (Figure 3 and Figure 7) (Hartmann et al., 2018). The induction of *UGT76B1* by *Pseudomonas* infection therefore suggests that it plays a major role in regulating NHP homeostasis, which seems to be critical to balance growth and defense in plants.

Although NHP level is higher in *ugt76b1* mutants, the increased accumulation of SA is most likely due to the reduced conversion of SA to SAG rather than the effect of NHP on the transcript levels of SA biosynthesis genes (Figure 4). In addition, the *FMO1*-3D mutant does not accumulate free SA to higher levels then the wild type and a lack of NHP does not affect the accumulation of SA in *fmo1*-1 plants (Koch et al., 2006; Bartsch et al., 2010). The increase of SA and NHP levels in *ugt76b1* mutants suggest that reduced turnover could be a critical mechanism for increasing the accumulation of SA as well as NHP (Figure S6). As there are three UGTs described to glycosylate SA, reduced accumulation of SAG could also hint for a deregulation mechanism in *ugt76b1*-1 plants towards the previously described SA UGTs, especially SAG-forming enzyme UGT74F1 (Dean and Delaney, 2008; George Thompson et al., 2017). Increased basal SGE level in *ugt76b1*-1 has already been addressed and connected to high basal *PR1* expression (von Saint Paul et al., 2011). However, after pathogen infiltration with *P.s.m*. transcript levels of *PR1* are similar in Col-0 and *ugt76b1*-1 (Figure S3). Furthermore, the transcript levels of *UGT74F2* coding for the SGE forming enzyme were similar in wild type and the *ugt76b1*-1 mutant. We conclude that the reported increase of SGE after infection of *ugt76b1*-1 is likely caused by the accumulation in UGT74F2s substrate SA (Figure 1). ILA was previously identified as substrate of UGT76B1, however, it was not identified as a molecular marker of infection with *Pseudomonas* in our non-targeted metabolite fingerprinting approach by UHPLC-HRMS (Supplemental Dataset 1). We observed neither ILA accumulation in *ugt76b1-*1, nor the respective glucoside in wild type plants after infection. Although there might be a chance that our workflow is not sufficient to detect these compounds *in vivo,* the intracellular concentration of ILA in the shoot was quantified to be approximately 2.5 ng per g dry weight and 7 ng per g dry weight for Col-0 and *ugt76b1*-1, respectively. Estimating a weight loss of at least 1:10 (*m/m*) between dry and f.w., the presented amounts of NHP are a multiple of ILA amounts in the shoot. Considering the determined K_M_ value of UGT76B1 for NHP in comparison with the one towards ILA presented earlier (472±97 μM) we consider ILA of minor importance for the observed enhanced resistance phenotype (Maksym et al., 2018). Most likely the enhanced resistance phenotype of *ugt76b1-*1 is therefore due to increased accumulation of NHP and SA.

The similar K_M_-values for NHP and SA suggest that UGT76B1 has a similar substrate specificity towards these two molecules. Additionally, the K_M_-value for SA determined in this work is similar to earlier reports (Noutoshi et al., 2012; Maksym et al., 2018). Nevertheless, NHP and SA differ in their absolute amount in infected leaf material (Figure 2a and 2b) to several orders of magnitude, suggesting that NHP is the more accessible, therefore, preferred substrate of UGT76B1. Although amino acid sequence comparison of UGT74F1 and UGT74F2 with UGT76B1 revealed only 26.96 % and 26.75 % sequence identity respectively, two critical residues for glycosylation (His20 and Asp109) in the putative active site are conserved among these UGTs (Figure S5) (George Thompson et al., 2017). Interestingly, we were not able to detect glycosylation of 4-OH-BA by UGT76B1 neither at the hydroxyl group nor at the carboxyl group. This suggests that a hydroxyl group in *ortho* or *meta* configuration adjacent to the carboxyl function is important for optimal binding of the ligand in the active side of UGT76B1.

From our transport experiments with D_9_-NHP, we conclude that NHP is not only a mobile signal, but can translocate from the apoplast to the cytosol and, rather than NHP-*O*Glc, is required for the establishment of SAR. This may be supported by an earlier study in which SAG was infiltrated into tobacco leaves (Hennig et al., 1993). Here, the authors showed that SAG was hydrolyzed in the apoplast to SA and that rather SA than SAG entered the cell. In addition, other studies support our notion that both NHP and SA are mobile between local and systemic tissue in *Arabidopsis* and tobacco (Yalpani et al., 1991; Chen et al., 2018; Lim et al., 2020). Nevertheless, it is still a matter of debate, as there was also evidence presented that SA is not the mobile signal for SAR (Vernooij et al., 1994b; Vernooij et al., 1994a). However, the formation of SAG and NHP-*O*Glc probably have a central role in inactivating SA and NHP as biological active molecules, as the dwarf phenotype of the corresponding mutant suggests (Figure 3c) (Noutoshi et al., 2012).

Based on the available data, we propose the following pathway for NHP-*O*Glc biosynthesis in Figure 8. First ALD1 converts L-lysine to P2C, which is then converted by SARD4 to Pip. Next, Pip is hydroxylated by FMO1 to NHP. In the last step, NHP is glycosylated by UGT76B1 to form NHP-*O*Glc. Together our data extend the NHP metabolic pathway down to NHP-*O*Glc and illustrates the major importance of UGT76B1 in metabolic regulation keeping defense and growth in balance.

**Figure 8:**
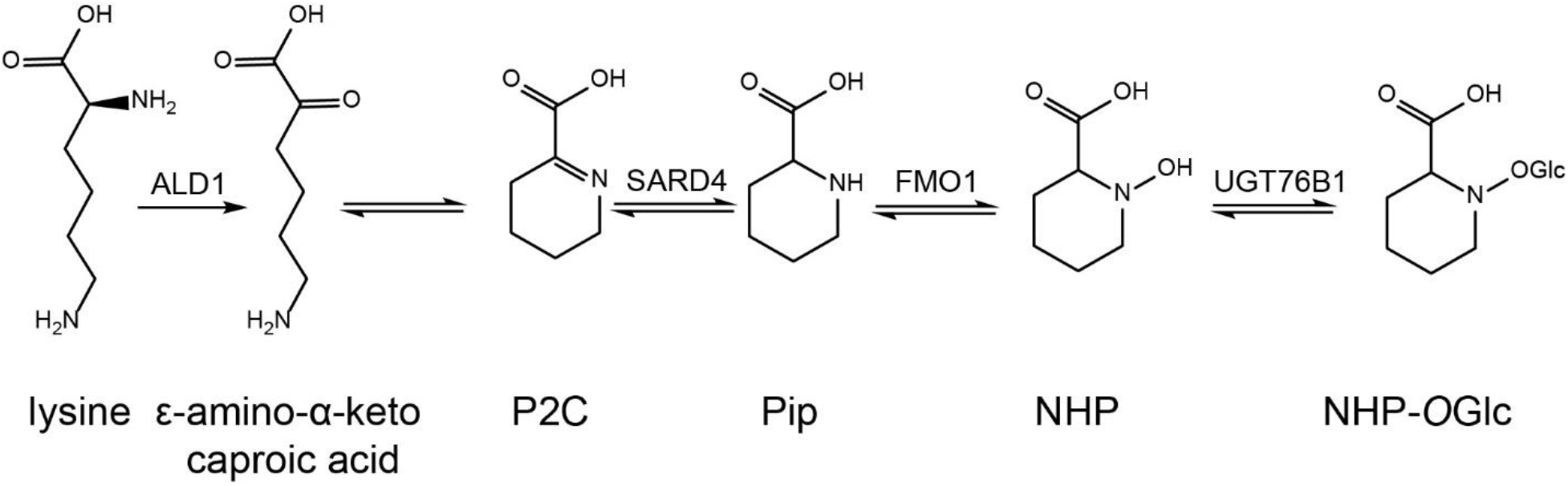
Biosynthesis of NHP-*O*Glc. The biosynthesis of NHP-*O*Glc starts from L-lysine, which is converted by ALD1 to *ε*-amino-α-keto caproic acid (Navarova et al., 2012; Song et al., 2004; Vogel-Adghough et al., 2013). The compound spontaneously cyclizes to Δ^1^-piperideine-2-carboxylic acid (P2C) and is reduced by SAR-deficient 4 (SARD4) to pipecolic acid (Pip) (Ding et al., 2016; Hartmann et al., 2017). FMO1 hydroxylates pipecolic acid to form NHP, the biological active pipecolate (Chen et al., 2018; Hartmann et al., 2018). In a last step NHP is glucosylated at the hydroxyl function to form NHP-*O*Glc.

## Material and Methods

### Plant material and growth conditions

Plants used for this work are all in *A. thaliana* Col-0 ecotype background. The *fmo1*-1 and *ugt76b1-*1 (SAIL_1171_A11) T-DNA insertion lines were obtained from NASC (University of Nottingham) and they were described previously. *ugt76b1-*3 and *ugt76b1-*4 are independent *ugt76b1-*1 deletion lines generated by CRISPR-Cas9 in Col-0 background, with original lab code as CRISPR UGT #5 and #17 respectively. Double mutant lines *fmo1-*1 *ugt76b1*-40 and *fmo1-*1 *ugt76b1*-104 were generated by crossing *ugt76b1*-1 with *fmo1*-1. In addition, a CRISPR deletion line of *UGT76B1* was generated in *fmo1*-1 background and referred to as *fmo1*-1 *ugt76b1*-5. The overexpression mutant *FMO1*-3D was described previously (Koch et al., 2006). Plants were grown for 4-6 weeks under short day conditions (8 hours light/18 hours dark cycle) with 100-120 μmol/m^2^ per s of light intensity at 80 % relative humidity unless specified.

### Construction of plasmids for *UGT76b1* gene editing and generation of deletion mutants

Three deletion lines *ugt76b1-*3, *ugt76b1-*4 and *ugt76b1-*5 *fmo1-*1 (original lab code CRISPR UGT #5, CRISPR UGT #17 and CRISPR UGT in *fmo1* #1) were generated by CRISPR/Cas9 system as described (Xing et al., 2014). Two single guide RNAs were designed to target *UGT76B1* genomic DNA to generate a ~1,000 bp deletion. The PCR fragment containing the guide RNA sequences were amplified from the pCBC-DT1T2 vector with primers 3G11340-BsFF0 and 3G11340-BsRR0 and subsequently inserted into the pHEE401 vector using the BsaI site. The derived plasmid was transformed into *E. coli* and later *Agrobacterium* by electroporation. Col-0 and *fmo1-*1 plants were transformed with the *Agrobacterium* carrying the plasmid by floral dipping. T_1_ plants were screened for deletion mutants by PCR with primers listed in Table S1. Homologous deletion mutants were obtained in T_2_ generation.

### Elicitation of defense response by UV-C and *P.s.m.* ES4326

Plants were treated for 20 min with UV-C radiation in a sterile bench (Telstar Bio-II-A, Azbil Telstar Technologies, Barcelona, Spain). The sterile bench was pretreated for additional 20 min prior to radiating the plants. Untreated control plants and the UV-C-treated plants were harvested 24 hours later. Infection of plants was conducted by infiltrating plant leaves with *P.s.m.* ES4326 at OD_600_ = 0.05 in 10 mM MgCl_2_, if not state otherwise, to induce defense. The bacteria were grown in LB medium with Rifampicin (50 μg/μl). In the D_9_-NHP tracking experiment, 82 μg/ml of chemically synthesized D9-NHP was added to the infiltration solution.

### Metabolite extraction

Leaves were harvested 24 hours post infection and frozen in liquid nitrogen. The samples were ground under liquid nitrogen using Retsch 200 MM (Retsch, Haan, Germany). Ground material was weighed and extracted after a modified methyl-*tert*-butyl ether (MTBE) extraction (Feussner and Feussner, 2019). When metabolite quantification was desired, deuterium labeled D_9_-NHP, D_6_-SA and isotopically labeled ^13^C-SAG was added prior to extraction. The labeled compound serve as reference throughout the analysis in quantitative matter.

### UPLC-nanoESI-QTRAP-MS-based metabolite quantification

Absolute quantification of NHP, NHP-*O*Glc, SA and SAG was performed corresponding to a method previously described (Herrfurth and Feussner, 2020), including the addition of 50 ng D_9_-NHP (kindly provided by Prof. Ulf Diederichsen, Goettingen, Germany), 10 ng D_4_-SA (C/D/N Isotopes Inc., Pointe-Claire, Canada) and 50 ng ^13^C_6_-SAG (kindly provided by Prof. Petr Karlovsky, Goettingen, Germany). Multiple reaction monitoring (MRM) transitions analyzed are shown in supplementary table 2. D_9_-NHP was synthesized as described previously (Hartmann et al., 2018). Synthesized NHP was characterized via tandem MS (MS/MS) fragmentation (Rekhter et al., 2019a). The fragmentation behavior underlying the MRM transitions of NHP-*O*Glc were analyzed after thin layer chromatographically purification of enzymatically produced NHP-*O*Glc using UGT76B1. As stationary phase a TLC silica gel 60(Merck KGaA, Darmstadt, Germany) was used in combination with butanol:water:acetic acid (4:1:1, *v/v/v*) as solvent system (Song, 2006). Purified NHP-*O*Glc was extracted from the silica gel with MTBE corresponding to the extraction procedure as described (Herrfurth and Feussner, 2020). Successful purification of enzymatically produced NHP-*O*Glc was checked via non-targeted UHPLC-HRMS. The quantification of the purified NHP-*O*Glc was performed by direct infusion-MS with respect to SAG (kindly provided by Prof. Petr Karlovsky, Goettingen, Germany).

### UHPLC-HRMS-based metabolite fingerprint analysis

Metabolites were extracted from 100 mg leaf material by two-phase extraction with MTBE, methanol and water according to Feussner and Feussner, 2019. Metabolite fingerprint analysis of the metabolites of the polar extraction phase was performed with the UHPLC1290 Infinity (Agilent Technologies) coupled to a HRMS instrument (6540 UHD Accurate-Mass Q-TOF, Agilent Technologies) with Agilent Dual Jet Stream Technology as electrospray ionization (ESI) source (Agilent Technologies). For chromatographic separation an ACQUITY HSS T3 column (2.1 × 100 mm, 1.8 μm particle size, Waters Corporation) was used with a flow rate of 500 μl/min at 40 °C. The solvent systems A (water, 0.1 % (*v/v*) formic acid) and B (acetonitrile, 0.1 % (*v/v*) formic acid) were used for the following gradient elution: 0 to 3 min: 1 % to 20 % B; 3 to 8 min: 20 % to 97 % B; 8 to 12 min: 100 % B; 12 to 15 min: 1 % B. The QTOF MS instrument was used in a range from *m/z* 50 to *m/z* 1700 with a detection frequency of 4 GHz, a capillary voltage of 3000 V, nozzle and fragmentor voltage of 200 V as well as 100 V, respectively. The sheath gas was set to 300 °C, and gas to 250 °C. The gas flow of drying gas was set to 8 l/min and sheath gas to 8 l/min, respectively. Data were acquired with Mass Hunter Acquisition B.03.01 (Agilent Technologies) in positive as well as ESI mode.

For data deconvolution the software Profinder B.08.02 (Agilent Technologies) was used. For further data processing, statistics, data mining and visualization the tools of the MarVis-Suite (Kaever et al. 2015, http://marvis.gobics.de/) was applied. Overall, 448 metabolite features (307 features from positive and 141 features from negative ESI mode) with a FDR < 0.005 were selected and clustered by means of one-dimensional self-organizing maps. The accurate mass information the metabolite features was used for metabolite annotation (KEGG, http://www.kegg.jp and BioCyc, http://biocyc.org, in-house database). The chemical structure of the indicated metabolites were confirmed by HRMS^2^ analyses (NHP: [M+H]^+^ 146.080, 128.070, 110.06, 100.076, 82.065, 70.065, 55.055 (Rekhter et al., 2019b); NHP-*O*Glc: [M+H]^+^ 308.132, 146.081, 128.0705, 110.06, 100.076, 82.062, 70.065, 55.055 (Rekhter et al., 2019b); SA: [M-H]^−^ 137.025, 93.035 (METLIN (https://metlin.scripps.edu/), MID3263); SAG: [M-H]^−^ 299.0719, 137.024, 93.035; Pip: [M+H]^+^ 130.086, 84.081, 70.065, 56.050 (Ding et al., 2016); 2HNG: [M-H]^−^ 216.051, 172.062, 128.072, 86.025 (Rekhter et al., 2019b) and SGE: [M-H]^−^ 299.078, 137.024, 93.035). The results were confirmed by two independent experiments with three biological replicates each.

### RNA extraction, Reverse Transcription and Quantitative Real-time PCR

Plants for gene expression assay were grown on soil under long-day (16 h light) condition. Three leaves of four-week-old plants (~50 mg) were collected for RNA extraction by EZ-10 Spin Column Plant RNA Miniprep Kit (Bio Basic Canada). RNAs were then reverse transcribed into cDNAs by OneScript Reverse Transcriptase (Applied Biological Materials Inc.). qPCR was performed with cDNAs using SYBR Premix Ex Taq™ II (Takara, Japan). For pathogen-induced gene expression assay, plants were grown under short-day (12h light) condition. Three leaves of four-six weeks old plants were infiltrated with *P.s.m.* ES4326 (OD_600_=0.001). Leaves were harvested 24 hpi and analyzed via the process as above. Primers for qPCR were listed in Table S1.

### Heterologous protein expression and purification

His-tagged UGT76B1 was purified via a combination of methods described recently (Maksym et al., 2018; Haroth et al., 2019). UGT76B1 (*AT3G11340,* GenBank Accession Number Q9C768.1) was amplified from total cDNA derived from infected leave tissue and cloned into pET28a vector (Merck, Darmstadt, Germany) using the BamHI and SalI restriction sites. The plasmid containing the *UGT76B1* gene was transformed into BL21 Star (DE3) cells (Thermo Fisher Scientific, Waltham, MA, USA) by heat shock. Cell cultures were grown in auto-induction medium (Studier, 2005) at 16 °C for 4 d. Cell pellets of 1 liter culture were resuspended in lysis buffer (50 mM Tris/HCl pH= 7.8, lysozyme, DNAseI and 0.1 mM PMSF). After homogenization, cells were disrupted by ultrasonication. Cleared lysate was obtained by centrifugation at 25000 xg for 45 min. The recombinant protein was purified from the cleared lysate using a combination of metal affinity chromatography using nickel-affinity (GE Healthcare, Chicago, IL, USA) and size exclusion chromatography using 16/600 Superdex 75 prep grade columns (GE Healthcare, Chicago, IL, USA).

### LC-MS based activity assay and *in vitro* kinetics

UGT76B1 recombinant protein was incubated with substrates NHP, SA and ILA for 30 min at 30 °C. The reaction was stopped by the addition of 20 % acetonitrile. Samples were analyzed using a 1290 Infinity UHPLC system coupled to a 6540 UHD Accurate-Mass Q-TOF (Agilent Technologies, USA) as previously described (Feussner and Feussner, 2019). Kinetic parameters of UGT76B1’s substrates NHP, SA and ILA were analyzed via UHPLC-HRMS. The reaction mixture contained 3.5 μg UGT76B1, 2 mM UDP-Glc (Merck KGaA) and 0-2.5 mM substrate. Before the incubation with UGT76B1, the initial amount of substrate was determined for analysis of substrate reduction. The reaction was incubated for 15 min at 30 °C and stopped by the addition of MeOH. The difference in signal intensity of substrate was plotted for each substrate and concentration. The Michaelis-Menten constant K_M_ was determined via Hill regression analysis using OriginPro8.5 (OriginLab Corporation, Northampton, MA, USA).

### Pathogen infection assay and SAR assay

Basal resistance against *H.a.* Noco 2 was tested by spay-inoculating two-week-old seedlings with spore solution (50,000 spores/mL). Inoculated seedlings were covered by a transparent lid and grown in a plant chamber with a relative humidity of ~80 %. Infection was scored 7 dpi by counting conidia spores with a hemocytometer.

Induction of SAR against *H.a.* Noco 2 was performed by infiltrating two full-grown leaves of three-week-old plants with *P.s.m.* ES4326 (OD_600_ = 0.001) or 10 mM MgCl_2_ (mock). Two days later, plants were sprayed with *H.a.* Noco 2 spore solution (50,000 spores/mL). Infection on distal leaves were scored 7 dpi as described previously (Ding et al., 2016).

Induction of SAR against *Pseudomonas* was tested by infiltrating *P.s.m.* ES4326 (OD_600_ = 0.001) or 10 mM MgCl_2_ (mock) on two leaves of four-week-old plants grown under short-day condition. Two days later, two distal leaves were challenged with *P.s.m.* ES4326 (OD_600_ = 0.001). Infection was scored both 0 dpi and 3 dpi by measuring the bacterial titer in the distal leaves.

### Structural prediction and ligand docking

The crystal structure of UGT74F2 (George Thompson et al., 2017), co-crystalized with SA-analogue 2-bromobenzoic acid, UDP, 3-*O*-β-D-glucopyranosyl-β-D-glucopyranose and β-D-glucose (PDB ID 5V2J) was used for structural prediction of UGT76B1. The structural prediction of UGT76B1 was done by PHYR2Protein (Kelley et al., 2015). NHP was fit into the electron density of SA-analogue 2-bromobenzoic acid using Coot (Emsley and Cowtan, 2004). Figures were created and distances were measured using PyMol (Schrödinger LLC, USA).

### Statistical analysis

Statistical analysis were performed using Origin Pro8.5 (OriginLab Corporation, Northampton, MA, USA).

## Supplemental Data

**Supplemental Dataset 1.** Non-targeted metabolite fingerprinting of Col-0, *fmo1*-1 and *ugt76b1*-1 after P.s.m. infection.

## Acknowledgments

We would like to acknowledge Brigitte Worbs for the chemical synthesis of the NHP and D_9_-labeled NHP standard. We thank Prof. Dr. Petr Karlovsky for kindly providing the SAG standard. LM and DR were supported by the Goettingen Graduate School for Neurosciences, Biophysics, and Molecular Biosciences (GGNB) at the Georg August University Goettingen. IF acknowledges funding from the Deutsche Forschungsgemeinschaft (DFG; GRK 2172-PRoTECT, INST 186/822-1 and ZUK 45/2010). YZ acknowledges funding from the NSERC Discovery Program. WH was supported by China Scholarship Council and NSERC-CREATE (PRoTECT).

## Author contributions

YZ and IF designed and supervised the study. Experimental research was conducted by LM, DR, WH, KF, HT and CH. LM, DR, WH, KF, CH, YZ and IF analyzed the data and wrote the manuscript.

## Supplemental Figures and Tables

**Figure S1.**
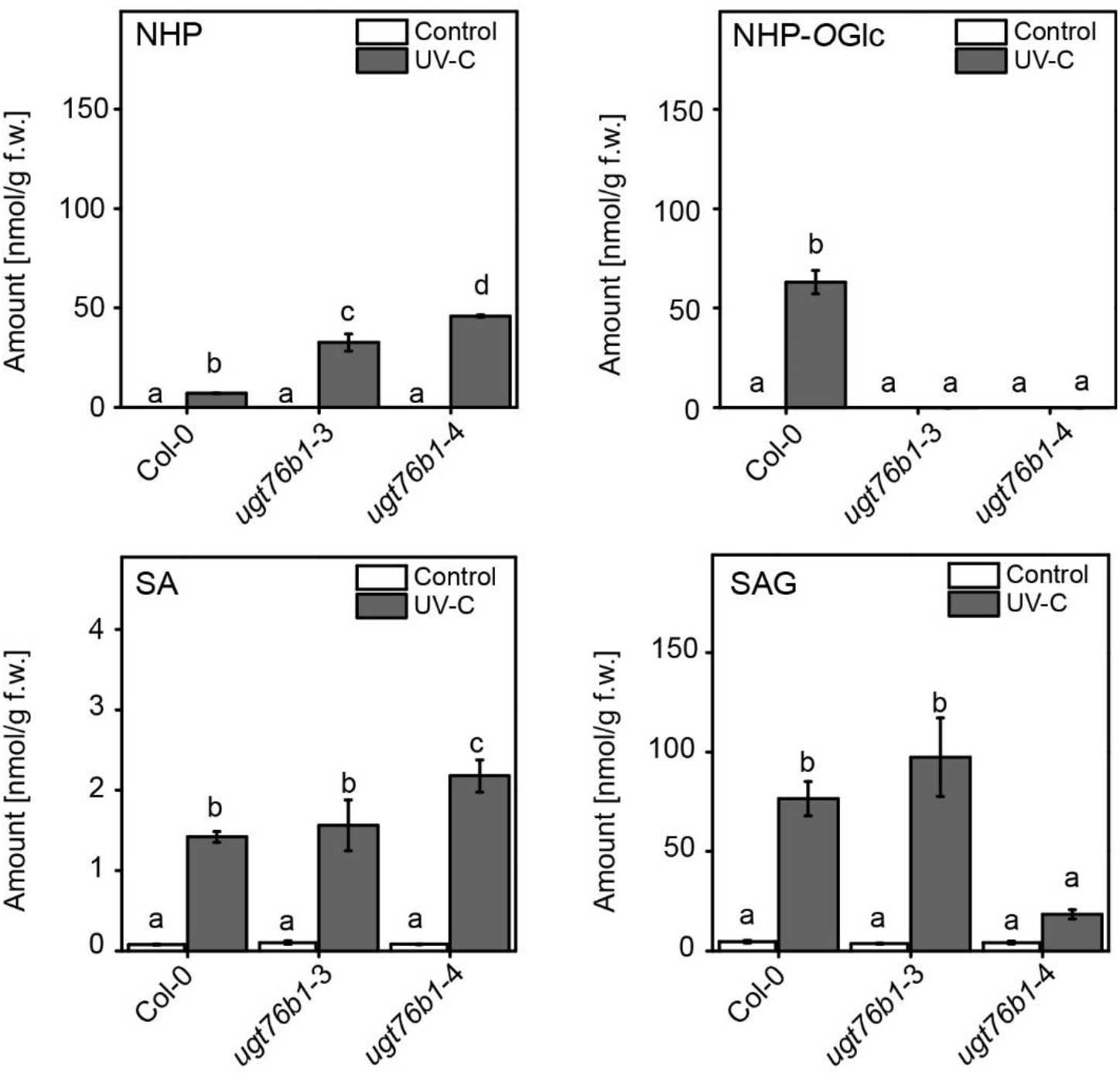
CRISPR deletion mutants of *UGT76B1* are unable to synthesized NHP-*O*Glc after UV-treatment. Absolute amounts of NHP, NHP-*O*Glc, SA and SAG were determined in wild type, *ugt76b1*-3 and *ugt76b1-*4 after UV-C treatment. Plants grown under long day conditions (16 hours light period), were treated for 20 min with UV-C or left untreated as control. 24 hours post treatment, leave material was harvested and analyzed using UPLC-nanoESI-QTRAP-MS. Letters indicate statistical differences (p < 0.05, one-way ANOVA; n=3 biological replicates).

**Figure S2.**
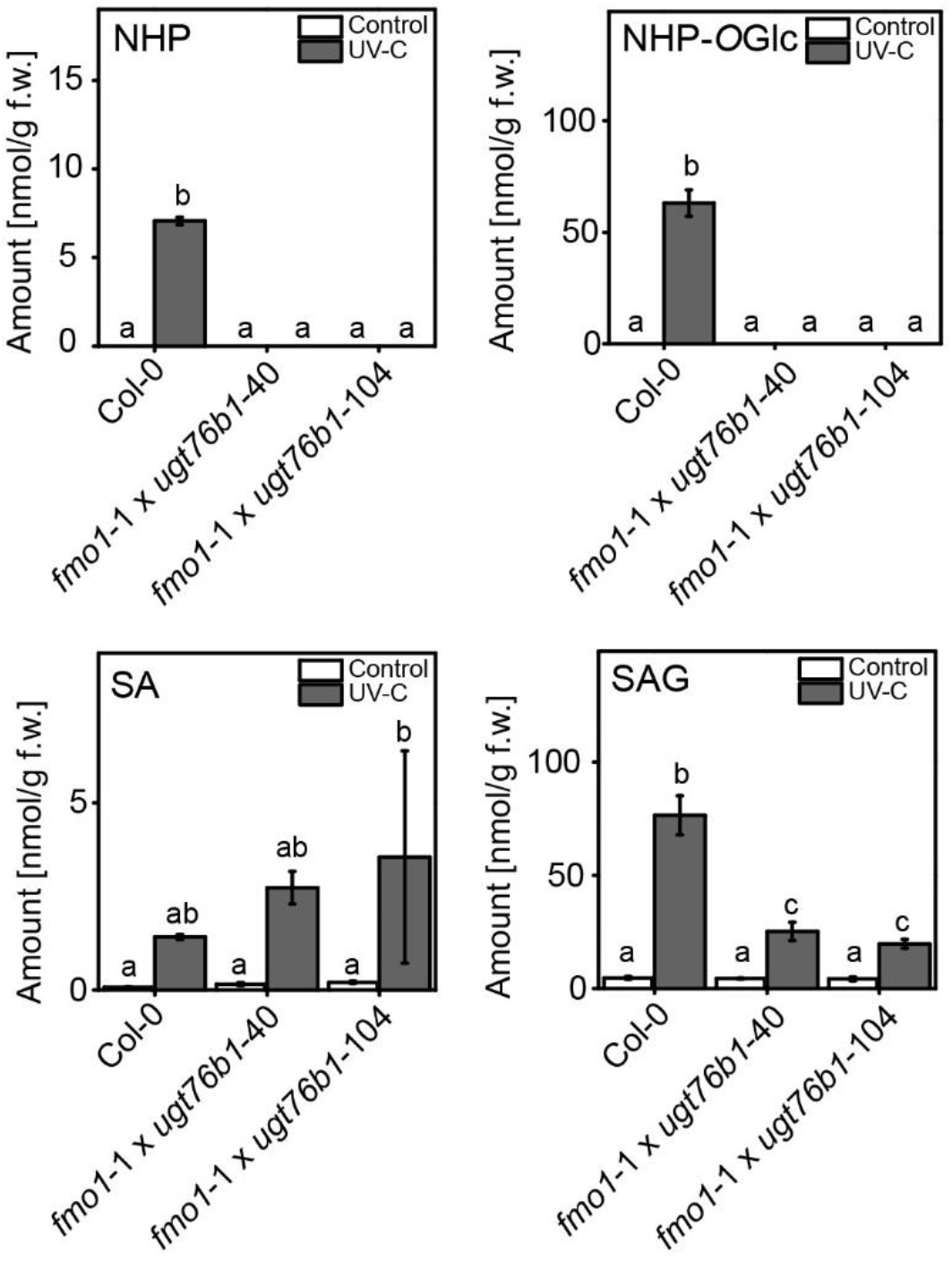
*fmo1*-1 *ugt76b1*-1 double loss-of-function mutant plants synthesize neither NHP nor NHP-*O*Glc after UV-treatment. Absolute amounts of NHP, NHP-*O*Glc, SA and SAG were determined in wild type and two independent *fmo1*-1 *ugt76b1* lines after UV-C treatment. Plants grown under long day conditions (16 hours light period), were treated for 20 min with UV-C or left untreated as control. 24 hours post treatment leave material was harvested and analyzed using UPLC-nanoESI-QTRAP-MS. Letters indicate statistical differences (p < 0.05, one-way ANOVA; n=3 biological replicates).

**Figure S3.**
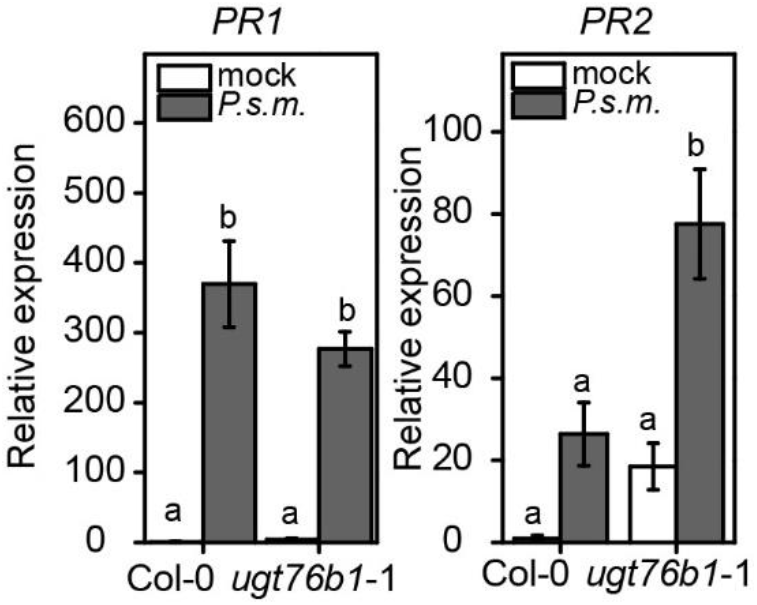
Transcripts levels of *PR1* and *PR2* after infection with *P.s.m*. in *ugt76b1* and wild type. Relative amount of transcripts of *PR1* and *PR1* was analyzed in wild type and *ugt76b1*-1 plants after infection with *P.s.m.* ES4326. Three leaves of 4-6 week-old plants were treated with *P.s.m.* ES4326 (OD_600_=0.001). Leaves were harvested 24 hours post infiltration and analyzed for the level of transcripts via quantitative RT-PCR. Letters indicate statistical differences (p < 0.05, one-way ANOVA; n=3 biological replicates).

**Figure S4.**
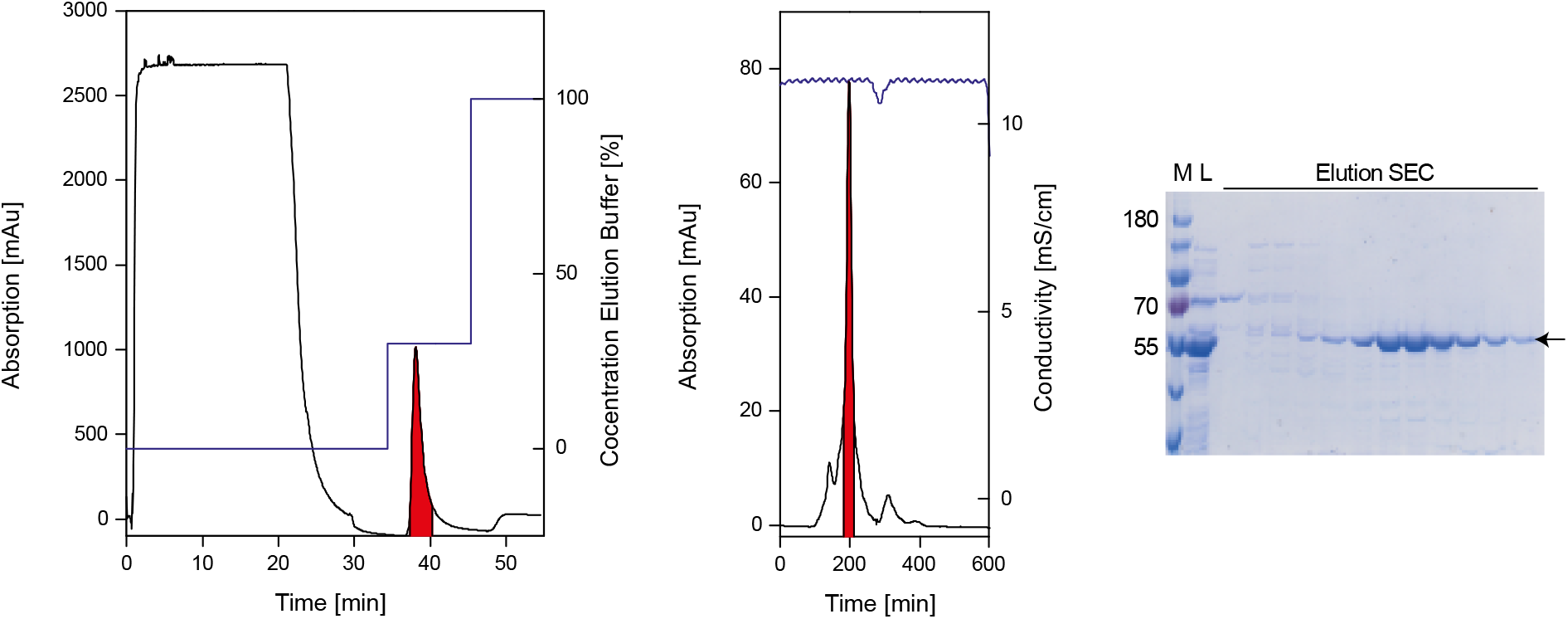
Purification of UGT76B1 heterologously expressed in *E. coli*. UGT76B1 fused with an N-terminal His-tag was heterologously expressed in *E. coli* BL21 Star (DE3) and purified via a combination of immobilized metal affinity chromatography (IMAC) and size exclusion chromatography (SEC). Chromatograms illustrate the absorption at 280 nm in milli absorption units (mAU) during protein elution. Secondary y-axes indicate the concentration of elution buffer in % for IMAC or the conductivity in mS/cm for SEC. Red areas represent corresponding signals to UGT76B1. The sodium dodecyl sulfate polyacrylamide gel electrophoresis (SDS-PAGE) shows the corresponding protein marker M, the load L (eluate IMAC) and the elution after SEC. The arrow indicates UGT76B1. The depicted purification is representative for at least three independent purification.

**Figure S5.**
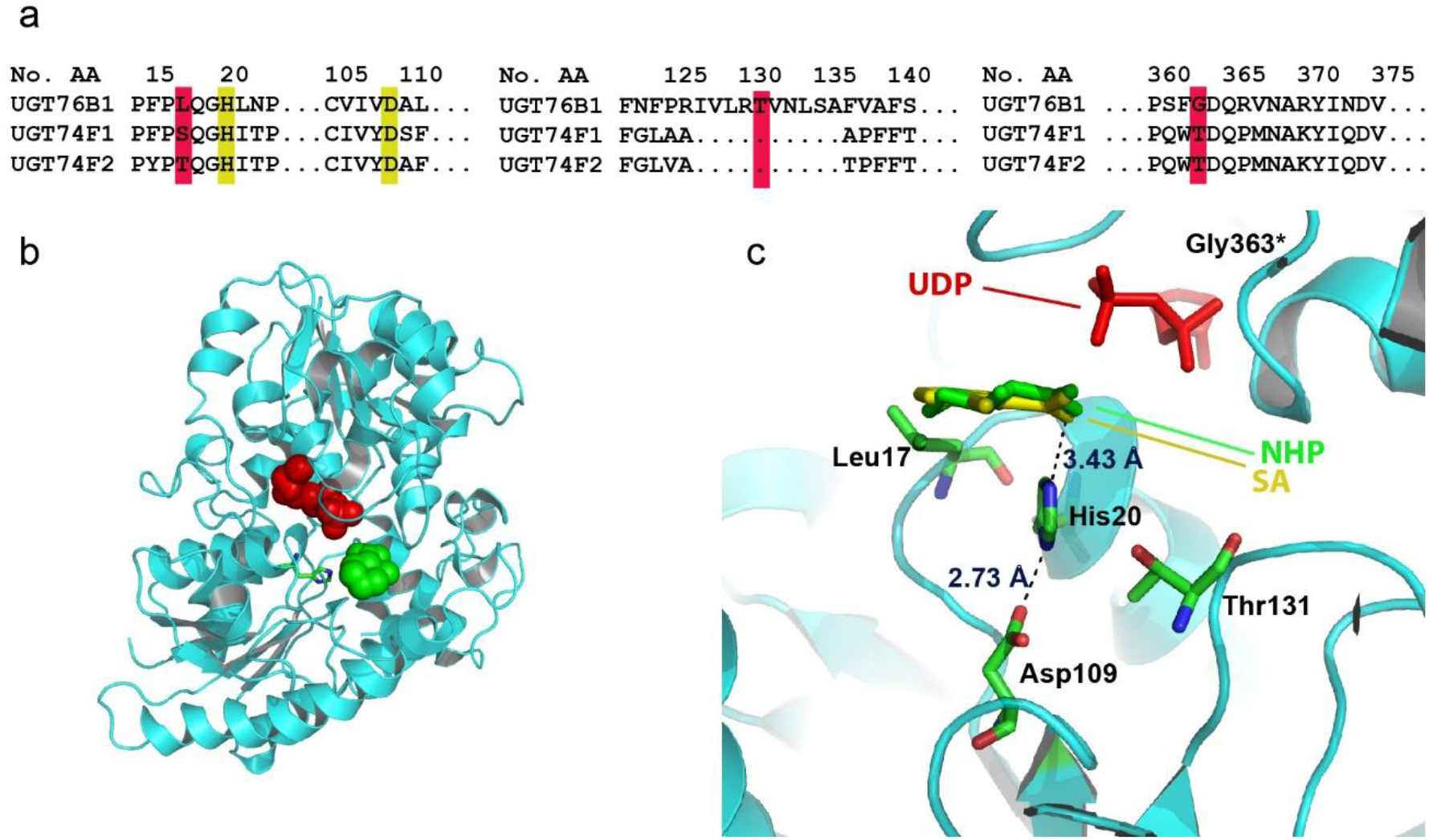
Modeling of NHP into the SA-analogues electron density in the predicted *in silico* UGT76B1 model. **(a)** Protein sequence alignment comparing UGT76B1, UGT74F1 and UGT74F2 towards the putative active site residues. Sequence identities are shown in yellow and miss matches in red. **(b)** Predicted model of UGT76B1 complexed with UDP and NHP using the deposited PDB structure 5V2J of UGT74F2 complexed with UDP and SA. UDP is show as balls in red and the modeled NHP is shown as balls in green. His20 is shown as sticks. **(c)** Amino acids histidine (His^20^), aspartate (Asp^109^) and putatively threonine (Thr^131^), which may form the proposed catalytic triad by George Thompson et al., 2017, are predicted to the active center and in close proximity to the substrate and each other in the UGT76B1 model prediction. The structural prediction of UGT76B1 was done by PHYR2Protein (Kelley et al., 2015). NHP was fit into the electron density of SA-analogue 2-bromobenzoic acid using Coot (Emsley and Cowtan, 2004). Figures were created using PyMol (Schrödinger LLC, USA).

**Figure S6.**
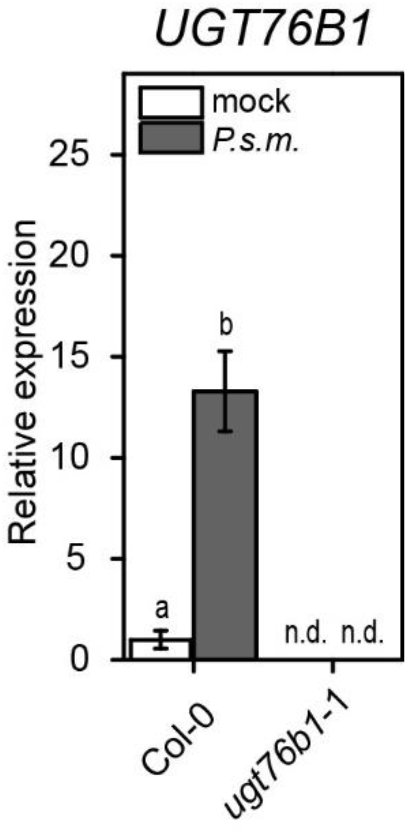
Transcripts of *UGT76B1* were not present in the mutant. Relative amount of transcripts of *UGT76B1* was analyzed in wild type and *ugt76b1*-1 plants after infection with *P.s.m.* ES4326. Three leaves of 4-6 week-old plants were treated with *P.s.m.* ES4326 (OD_600_=0.001). Leaves were harvested 24 hours post infiltration and analyzed for the level of transcripts via quantitative RT-PCR. Letters indicate statistical differences (p < 0.05, one-way ANOVA; n=3 biological replicates).

**Table S1.**
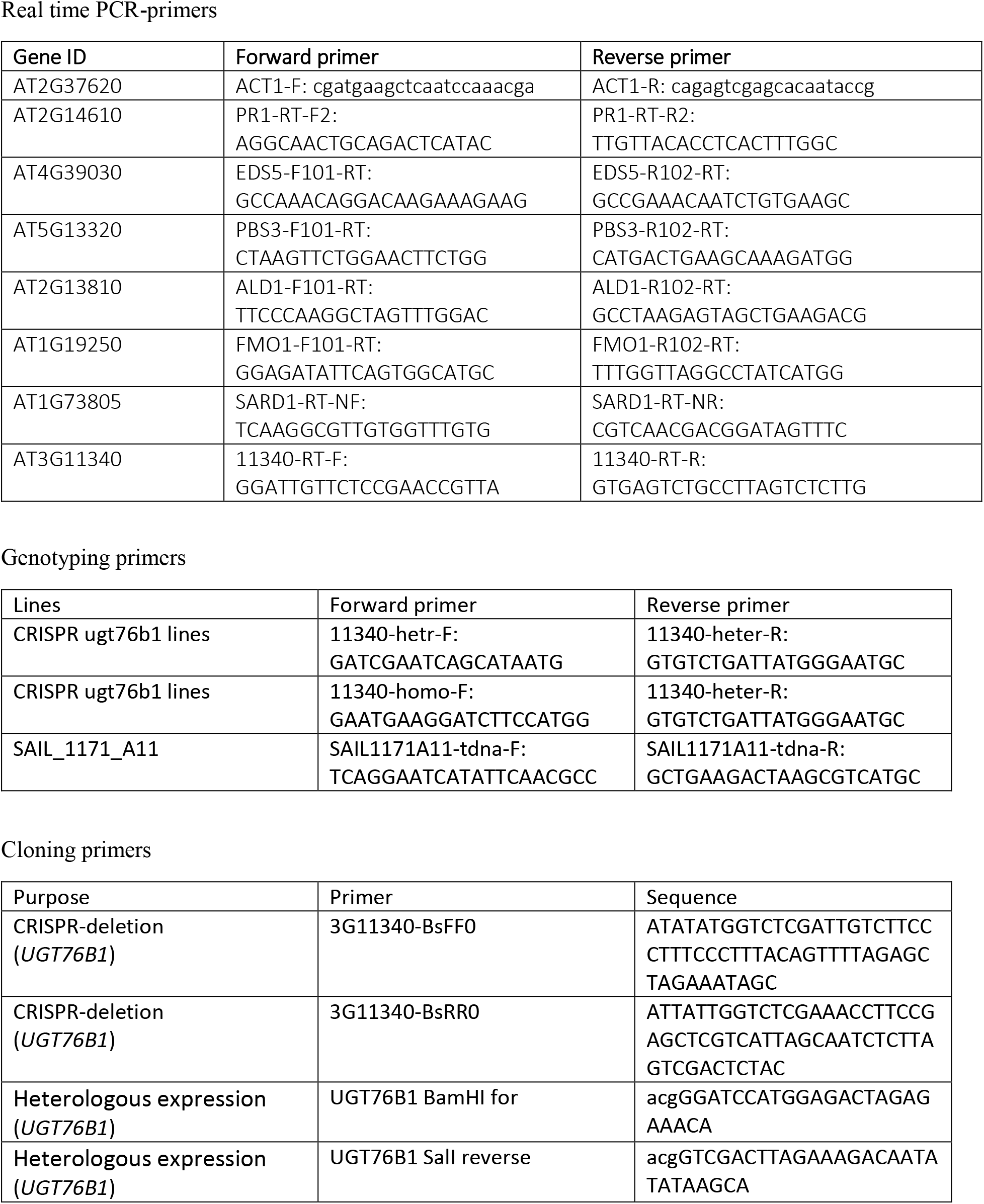
List of primers used in this work. Information is divided by primer application for quantitative PCR analysis, genotyping, and cloning.

**Table S2.** Multiple reaction monitoring parameters for absolute quantification of analytes. For the presented quantitative plant hormone data we established a multiple reaction monitoring analysis of seven additional analytes to the ones published before (Herrfurth and Feussner, 2020). Information are divided for mass spectrometric analysis after negative ionization or positive ionization. Q1 (precursor ion), Q3 (product ion) and the retention time (RT) of each analyte are shown, respectively. Furthermore, the declustering potential (DP), entrance potential (EP), collision energy (CE) and the cell exit potential (CXP) of each compound are provided.

| **Q1**<br>**(m/z)** | **Q3**<br>**(m/z)** | **RT**<br>**(min)** | **Analyte** | **DP**<br>**(eV)** | **EP**<br>**(eV)** | **CE**<br>**(eV)** | **CXP**<br>**(eV)** |
| --- | --- | --- | --- | --- | --- | --- | --- |
| 137 | 93 | 2 | SA | -25 | -6 | -20 | -10 |
| 141 | 97 | 3 | D<sub>4</sub>-SA | -25 | -6 | -22 | -6 |
| 144 | 82 | 0.7 | NHP | -60 | -8 | -15 | -13 |
| 153 | 90 | 0.7 | D<sub>9</sub>-NHP | -60 | -8 | -15 | -13 |
| 299 | 137 | 1 | SAG | -30 | -4 | -18 | -2 |
| 305 | 137 | 1 | <sup>13</sup>C<sub>6</sub>-SAG | -30 | -4 | -18 | -2 |
| 306 | 89 | 0.9 | NHP-OGlc | -65 | -4 | -18 | -13 |

positive nanoESI
| **Q1**<br>**(m/z)** | **Q3**<br>**(m/z)** | **RT**<br>**(min)** | **Analyte** | **DP**<br>**(eV)** | **EP**<br>**(eV)** | **CE**<br>**(eV)** | **CXP**<br>**(eV)** |
| --- | --- | --- | --- | --- | --- | --- | --- |
| 130 | 84 | 0.7 | PIP | 90 | 8 | 22 | 4 |
| 139 | 93 | 0.7 | D<sub>9</sub>-PIP | 90 | 8 | 22 | 4 |

